# Agronomic treatments combined with embryo rescue for rapid generation advancement in tomato speed breeding

**DOI:** 10.1101/2023.02.28.530438

**Authors:** Esther Gimeno-Páez, Jaime Prohens, María Moreno-Cerveró, Ana de Luis-Margarit, María José Díez, Pietro Gramazio

## Abstract

Unlike other major crops, little research has been performed on tomato for reducing generation time for speed breeding. We evaluated several agronomic treatments for reducing the generation time of tomato in the M82 (determinate) and Moneymaker (indeterminate) varieties and evaluated the best combination in conjunction with embryo rescue. In a first experiment under the autumn cycle, five container sizes, from 0.2 1 (XS) to 6 1 (XL), were evaluated. We found that plants from the XL containers exhibited better development and required less time from sowing to anthesis (DSA) and for anthesis to fruit ripening (DAR). In a second experiment, using XL containers in the autumn-winter cycle, we evaluated cold priming at the cotyledonary stage, water stress, P supplementation, and K supplementation on generation time. We found that, compared to the control, cold priming significantly reduced the number of leaves and plant height to first inflorescence as well as DSA (2.7 d), while K supplementation reduced DAR (8.8 d). No effects of these treatments were observed for other growth of physiological traits. In a third experiment with XL containers in the spring-summer cycle, the combination of cold priming plus K supplementation was tested, confirming the significant effect of the combination on generation time (2.9 d for DSA and 3.9 d for DAR). Embryo rescue during the cell expansion cycle (average of 22.0 d and 23.3 d after anthesis for M82 and Moneymaker, respectively) allowed shortening the generation time by 8.7 d in M82 and 11.6 d in Moneymaker compared to the *in planta* fruit ripening. The combination of agronomic treatments with embryo rescue can make an effective contribution to increase the number of generations per year for speed breeding in tomato from the current three to four.

## Introduction

Rapid generation advancement is one of the cornerstones of speed breeding. By reducing the generation time, the length of breeding programs is shortened, which allows speeding up the development of new varieties addressing the demands of consumers and the pressing challenges posed by climate change and the need for more sustainable agriculture (Wanga et al., 2021). One of the approaches for reducing the generation time is the modification of agronomic techniques, which are known to affect the generation time in many crops (Samantara et al., 2022). For example, embryo rescue, apart from being used for a long time to achieve hybrids in distant crosses (Sharma et al., 1996), can also be used to significantly shorten generation time in intraspecific crosses since it is not essential obtaining physiologically mature seeds to obtain the subsequent generation (Ghosh et al., 2018; Wanga et al., 2021). In crops amenable to the production of doubled haploids, homozygosis can be obtained in a single generation, which can shorten considerably the length of breeding programs when pure lines or fixation are needed (Wanga et al., 2021).

Despite its economic importance, compared to major cereal and legume crops (Gosal and Wani, 2020; Wanga et al., 2021; Samantara et al., 2022), tomato lags behind in the development of speed breeding techniques. For example, while for crops such as wheat, barley, oat, chickpea, pea, grass pea, canola, or quinoa there are specific speed breeding protocols (Ghosh et al., 2018), such framework has not been developed for tomato. In addition, tomato is highly recalcitrant to haploid induction and the development of efficient and genotype-independent doubled haploid protocols have failed so far (Hooghvorst and Nogués, 2021). Although immature seed culture and embryo rescue have proven very promising for rapid generation advancement in tomato (Bhattarai et al., 2009; Geboloğlu et al., 2011) their use has not been integrated with agronomic techniques easily applicable by breeding companies, such as the use of different container sizes, temperature treatments, irrigation, or fertilization treatments that may affect traits relevant for generation time such as flowering earliness or ripening.

Speed breeding requires cultivation all year round. When cultivation, as usually done by breeding companies, is performed under greenhouses, the time of the year in which the plants are cultivated affects the generation time. In this way, the cycle of cultivation is known to affect the length of the generation cycle of the tomato (Martín-Closas et al., 2009), with faster development under the higher light intensities and temperatures, as well as long photoperiods, of the spring-summer cycle.

Tomato can be grown in the soil or containers with a substrate. Cultivation in containers is often preferred in speed breeding (Ghosh et al., 2018). Container size influences the yield of tomato and, in general, the larger the pot size the higher the yield (Şirin and Sevgican, 1999). However, for the rapid advancement of generations yield is not an issue and using reduced pot sizes in speed breeding has the advantage of being able to grow more plants in less space. Little information is available on the effect of pot size on traits related to generation time in tomato. In this way, Ruff et al. (1987) compared tomato cultivation in 0.45 l and 13.5 l pots and found that plants in the small pots had a delay of around three days in anthesis and a slight delay in fruit maturation. However, they just used two very different container sizes and many more are available for tomato cultivation. In other crops such as cereals, small pot sizes are used to reduce generation time (Zheng et al., 2013; Ferrie and Polowick, 2020). For example, cotton growing in 2 l pots resulted in earlier flowering in comparison to 10 l pots (Carmi, 1986).

Furthermore, specific temperature treatments during sensitive periods can affect flowering in tomato. One of the first works, performed 70 years ago, reported that applying different periods of cold temperatures (14 °C) after cotyledon expansion affected the timing of initiation of flowering (Lewis, 1953). Subsequently, Calvert (1957) found that applying low temperatures (10-15 °C) for 9 d during the sensitive phase (after cotyledon expansion) allowed an early flowering response, as measured by a reduced number of leaves until the appearance of the first inflorescence, suggesting that cold treatments after cotyledon expansion could be used for reducing the generation time in tomato.

Drought is known to induce flowering in many species (Takeno, 2016) and is used in speed breeding in several crops (Wanga et al., 2020). In tomato, some works indicate that water deficit results in early flowering (Wudiri and Henderson, 1985; Chong et al., 2022), while in others, drought delays flowering and maturation in some genotypes (Martínez-Cuenca et al., 2020). In some genotypes, like the non-ripening mutant *nor*, drought induces ripening (Arad and Mizhari, 1983). However, the use of water deficit as a potential tool to shorten the generation time in tomato remains to be explored.

Modification of the macrominerals supplied with the fertilization may have an impact on traits related to generation time, such as flowering earliness and fruit ripening. In this way, in rice, it has been observed that increases in N fertilization delay flowering, while increases in P and K advance it (Ye et al., 2019). In wheat, it has been found that increasing ten times the concentration of KH_2_PO_4_ advanced flowering and allowed a shortening of the generation cycle in an in vitro protocol for speed breeding (Yao et al., 2017). Besford and Maw (1975) found that flowering time in tomato was advanced at high doses of K, while Dieleman and Heuvelink (1992) report that the number of leaves until the first inflorescence decreases at high levels of fertilization. Therefore, the evidence available suggests that P and K supplementation might be useful for reducing the different phases of generation time in tomato.

Aside from agronomic techniques, embryo rescue is also of interest for rapid generation advancement in tomato. Embryo rescue has been extensively used in tomato for interspecific hybridization since a long time ago (Kalloo, 1991; Picó et al., 2002). This has allowed the introgression of genes from tomato wild relatives with which sexual hybridization with the cultivated tomato is challenging or unfeasible (Díez and Nuez, 2008). However, embryo rescue has also been proposed as an efficient tool for rapid generation advancement in tomato breeding programs. In this way, Bhattarai et al. (2009) found that growing immature seeds at stages as early as 10 days (young heart stage embryos) after pollination resulted in a considerable reduction of the generation, facilitating the rapid advancement of generations compared to the standard seed-to-seed procedure (72 d vs. 132 d). However, at such early stages, embryos had lower germination and regeneration than those of more advanced stages. Other works, such as those of Demirel and Seniz (1997) or Geboloğlu et al. (2011), have found that later stages (from 20 to 30 d after pollination) are the most appropriate for the rapid advancement of generations in tomato. In this way, Geboloğlu et al. (2011) found that harvesting fruits at 20-24 d after pollination allowed a reduction of generation time of over one month when compared to the seed-to-seed control.

Given that, so far there have been no comprehensive investigations on reducing the generation time in tomato by combining different treatments. In this study, we present the experiments performed on environmental, physiological and tissue culture treatments for rapid generation advancement, either for backcrossing or rapid homozygotization in tomato. Among the treatments evaluated, we have performed a series of step-wise experiments devised to find a combination of agronomic techniques easy to adopt by breeding companies, which combined with embryo rescue can contribute to a significant reduction of generation time for speed breeding in tomato. The experiments are performed on indeterminate (‘Moneymaker’) and determinate (‘M82’) varieties widely used for research and breeding (Chaudhary et al., 2019). The resulting protocol/s may be combined with conventional or biotechnological (e.g., genetic transformation, gene editing, or transient expression) genetic approaches (Bauchet et al., 2017; Soyk et al., 2017; Adkar-Purushothama et al., 2018; Wang et al., 2019) aimed at reducing generation time in tomato.

## Materials and methods

### Plant materials and growing conditions

The tomato varieties ‘M82’ (determinate) and ‘Moneymaker’ (indeterminate) were used. For germination, seeds were sown in Petri dishes (9.0 x 2.5 cm) on a layer of embedded hydrophilic cotton covered by filter paper and placed in a growth chamber with a 16 h light / 18 h dark photoperiod at 25 °C (light) / 18 °C (dark) temperature. Light was provided by GRO-LUXF36W/GRO (Sylvania, Danvers, MA, USA) fluorescent tubes. Once the roots and cotyledon emerged, each seedling was transferred to a 0.2 l pot container filled with Neuhaus Huminsubstrat N3 growing substrate (Klassmann-Deilmann, Geeste, Germany), which is made of sphagnum frozen black peat and sphagnum white peat (organic matter content of 85%, pH of 6, conductivity of 35 mS/m, and water retention of 75%) enriched with 1 kg/m^3^ of a 14 N – 10 P_2_O_5_ – 18 K_2_O fertilizer. Plantlets were kept in the same growth chamber of germination until they reached the three true leaves stage. At this stage, depending on the experiment, they were either kept in the same container or transplanted to other larger containers for greenhouse evaluation. Containers were filled with the same Neuhaus Huminsubstrat N3 growing substrate.

Plants were grown on benches in a glasshouse at the Universitat Politècnica de València with climate control (heating started at temperatures below 15 °C and cooling at temperatures above 27 °C). For the experiment involving different container sizes (Experiment 1; EX1) plants were placed on top of concrete benches at a distance of at least 30 cm between individual plants. For the rest of the experiments (Experiments 2 and 3; EX2 and EX3), which involved only XL containers, plants were spaced 50 cm apart on benches with 115 cm between bench centers. Plants were watered manually every 1-3 days depending on the demands of the plants, which were determined by the stage of development and season. Plants from EX1 (different container sizes) were trained with bamboo canes or wood sticks, while those of the EX2 and EX3 (different agronomic treatments) were trained with vertical strings. No fertilization was provided in addition to the nutrients present in the substrate in EX1, while for EX2 and EX3 10 g per plant of a 14 N - 7 P_2_O_5_ - 17 K_2_O (+ 2 MgO) fertilizer (Nitrofoska 14, Eurochem Antwerpen NV, Antwerp, Belgium) were supplied to all plants as dressing fertilization 50 d after transplant. Some of the treatments of EX2 and EX3 involved extra P or K fertilization. Details are provided below in the “Treatments” subsection.

### Agronomic treatments

Three experiments (EX1-EX3) were performed involving agronomic treatments (Figure 1). EX1 was aimed at finding the best container size for rapid generation advancement. For this, five container sizes were evaluated: 0.2 1 (XS), 0.45 1 (S), 0.8 1 (M), 1.3 1 (L) and 6 1 (XL). Seeds were germinated on 24 August 2021 (autumn cycle) and 10 plants per combination of variety and container size were used. Plants were distributed according to a completely randomized block design. The container size (XL) that allowed the fastest generation advancement was used for subsequent experiments (EX2 and EX3).

**Figure 1.**
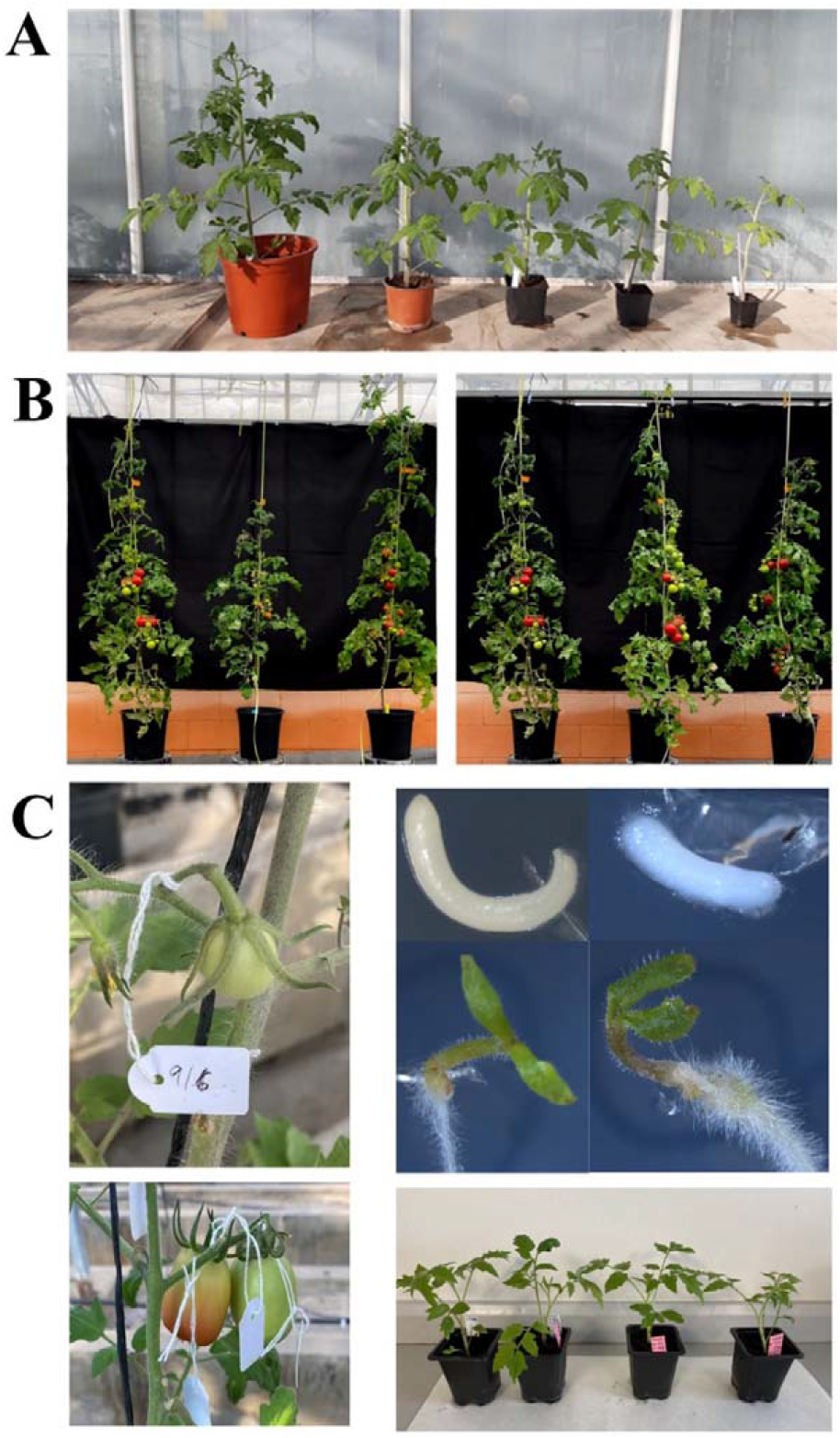
Treatments comparison for the three experiments performed in this study for reducing generation time in tomato. A) Experiment 1. Effects of container size. From left to right: 61 (XL), 0.8 1 (M), 0.45 1 (S), 0.21 (XS). B) Experiment 2. Effects of five treatments. From left to right: comparison of Control, Water Stress, Potassium treatment (left), Control, Phosphorus, and Cold Priming treatment (right) on Moneymaker. C) Experiment 3. Effect of Cold priming plus K supplementation and embryo rescue. Selfing and natural ripening fruits (left) versus embryo rescue (right).

In EX2, five treatments were compared: Control (C), Cold priming (CP), Water stress (WS), phosphorus supplementation (P), and potassium supplementation (K). For the cold priming treatment, once the cotyledon was fully expanded, plants were placed in a growth chamber with the same lighting photoperiod and conditions that the control, except that they were subjected to a constant temperature of 14 °C for eight days. After this priming period, plants from the cold treatment were moved to the control growth chamber with the rest of the plantlets from the other treatments. The water stress treatment consisted of reducing irrigation to half of the supply of the control, which was watered to field capacity. The water amounts to be supplied to the control were determined by measuring the substrate humidity with a WET-2 Sensor (Delta-T Devices, Cambridge, UK) and calculating the quantity of water required to reach field capacity. For the P and K supplementation treatments, each 6 l (XL) container was supplemented with 30 g of single superphosphate (18% P2O5; Fuentes Fertilizantes S.L.U., Totana, Spain) for the P supplementation or with 20 g of potassium sulfate (50% K_2_O; Antonio Tarazona S.L.U., Silla, Spain) for the K supplementation. Half of the amount of P or K supplementation was administered as dressing fertilization one week after transplant to the 6 l (XL) containers, while the other half at the start of the fruit set. For EX2, seeds were germinated on 8 October 2021 (autumn-winter cycle) and 10 plants per combination of variety and treatment were used. Plants were distributed on concrete benches according to a completely randomized block design.

In EX3, the best two individual treatments that allowed a significant reduction in generation time (cold priming and K supplementation) in EX2 were combined and compared to the Control. Treatments of cold priming and K supplementation were performed as in EX2. Seeds were put to germinate on 22 April 2022 (spring-summer cycle) and 20 plants per combination of variety and treatment were used. Ten of the plants of each combination of variety and treatment were randomly allocated for *in planta* ripening of the fruits (as in EX1 and EX2), while the other ten were left for embryo rescue. A completely randomized block design was used.

### Traits measured

The following morphological traits were evaluated in EX1 and EX2: number of leaves until the first inflorescence, stem diameter, plant height to the first inflorescence (cm), and distance between internodes (cm). Also, in EX1 and EX2 the chlorophyll index, anthocyanins index, flavonoids index, and Nitrogen Balance index (NBI) were taken with a Dualex-A optical sensor (Dualex Scientific^®^ (Force-A, Orsay, France). Dualex-A data were measured for the adaxial and abaxial sides of three young developed leaves per plant.

The time elapsed (d) from sowing to anthesis of the first flower (DSA) and from flower anthesis to first ripe fruit (red ripe stage; DAR) were counted for all plants in the three experiments, except the time from flower anthesis to first ripe fruit in the plants of EX3 devoted to embryo rescue. Instead, for the plants used for embryo rescue the time between anthesis and the first acclimatized plant with three true leaves (DA3L) was counted. In order to compare with the plants of EX3 in which the fruits ripened on plants, seeds of these latter fruits were germinated, and the time required for obtaining plants with three true leaves was counted (DS3L). For comparison of the time elapsed between anthesis and fruit ripening (DAR) of plants of EX3 in which fruits ripened on plants with those in which embryo rescue was applied, an equivalent to DAR (eDAR) was calculated for embryo rescue plants as eDAR=DA3L-DS3L.

### Embryo rescue

Immature fruits of plants from EX3 were harvested during the cell expansion phase (Table 1) at a stage considered appropriate for recovering torpedo and pre-cotyledonar embryos (Picó et al., 2002). After harvest, fruits were brought to the laboratory and surface sterilized with ethanol (96%) for 30 s in an AH-100 laminar flow cabinet (Telstar, Terrasa, Spain). Fruits were opened under sterile conditions in the same laminar flow cabinet and immature seeds were extracted and sterilized using a 1% dilution of commercial bleach (4% sodium hypochlorite) for 10 min (with two drops of Tween20) and rinsed three times with sterile distilled water for 1 minute. The immature seeds were dissected under a stereomicroscope Leica S8 APO (Leica Microsystems CMS GmbH, Wetzlar, Germany) at a magnification of 10x using sterilized dissection needles. Embryos were carefully excised and cultured in Petri dishes (9.0 x 2.5 cm) with the culture medium. The Petri dishes were sealed with Parafilm M (Amcor, Zurich, Switzerland) and moved to a growth chamber with a 16 h light / 18 h dark photoperiod at 25 °C (light) / 18 °C (dark) temperature. The lighting was provided by GRO-LUXF36W/GRO (Sylvania) fluorescent tubes.

**Table 1.** Average (±standard deviation) for the harvesting time and fruit length, width and weight of the fruits of M82 and Moneymaker used for embryo rescue (Experiment 3; EX3).

| Fruit characteristics | M82 | Moneymaker |
| --- | --- | --- |
| Harvesting time (days after anthesis) | 22.0±5.7 | 23.3±6.8 |
| Fruit length (mm) | 28.9±2.8 | 25.6±2.7 |
| Fruit width (mm) | 24.7±2.7 | 23.6±1.9 |
| Fruit weight (g) | 8.5±1.6 | 8.1±1.4 |

The culture medium for the incubation of rescued embryos consisted of 4.4 g/l of Murashige-Skoog salts, 30 g/l sucrose and, 7 g/L Gelrite™. All components were purchased from Duchefa Chemie (Harlem, The Netherlands). pH of the medium was adjusted to 5.9. The medium was sterilized by autoclaving at 121 °C for 20 min.

Once embryos developed cotyledons and root they were transferred to 0.87 l Microbox containers O118/120+OD118/120 (SAC O_2_, Deinze, Belgium) with the same in vitro MS medium until two leaves stage were reached. Then, they were removed from in vitro culture and were transferred to 0.2 l pots containing Huminsubstrat N3 growing substrate and covered with perforated plastic glasses to prevent dehydration and maintained in the same climatic chamber used for seed germination and growth of plantlets until they developed three true leaves.

### Statistical analysis

For each of the experiments, morphological data, Dualex-A indexes and times elapsed from sowing to anthesis (DSA) or from anthesis to fruit ripening (DAR or eDAR) were subjected to multifactorial ANOVA for the evaluation of the main effects of variety and container size (CS) or treatment (T) effects, as well as their respective double interactions (V x CS and V x T). Block effect was also calculated in order to reduce residual variation. Significance of differences among different levels of the main effects, as well as among combinations of main factors where interaction was significant (p<0.05), were evaluated using Duncan multiple range tests at p<0.05. All statistical analyses were conducted using the Statgraphics Centurion XVIII (v. 18.1.13) software (Statgraphics Technologies Inc., The Plains, VA, USA).

## Results

### Effects of container size on generation time (Experiment 1; EX1)

The ANOVA revealed significant (p<0.05) effects of the variety (V) and container size (CS) for all traits measured, except for the time from anthesis to ripening in the case of variety and internode length in the case of container size (Table 2). Interactions V x CS were non-significant, except for the number of leaves to the first inflorescence, plant height to first inflorescence and time from anthesis to ripening (Table 2).

**Table 2.**
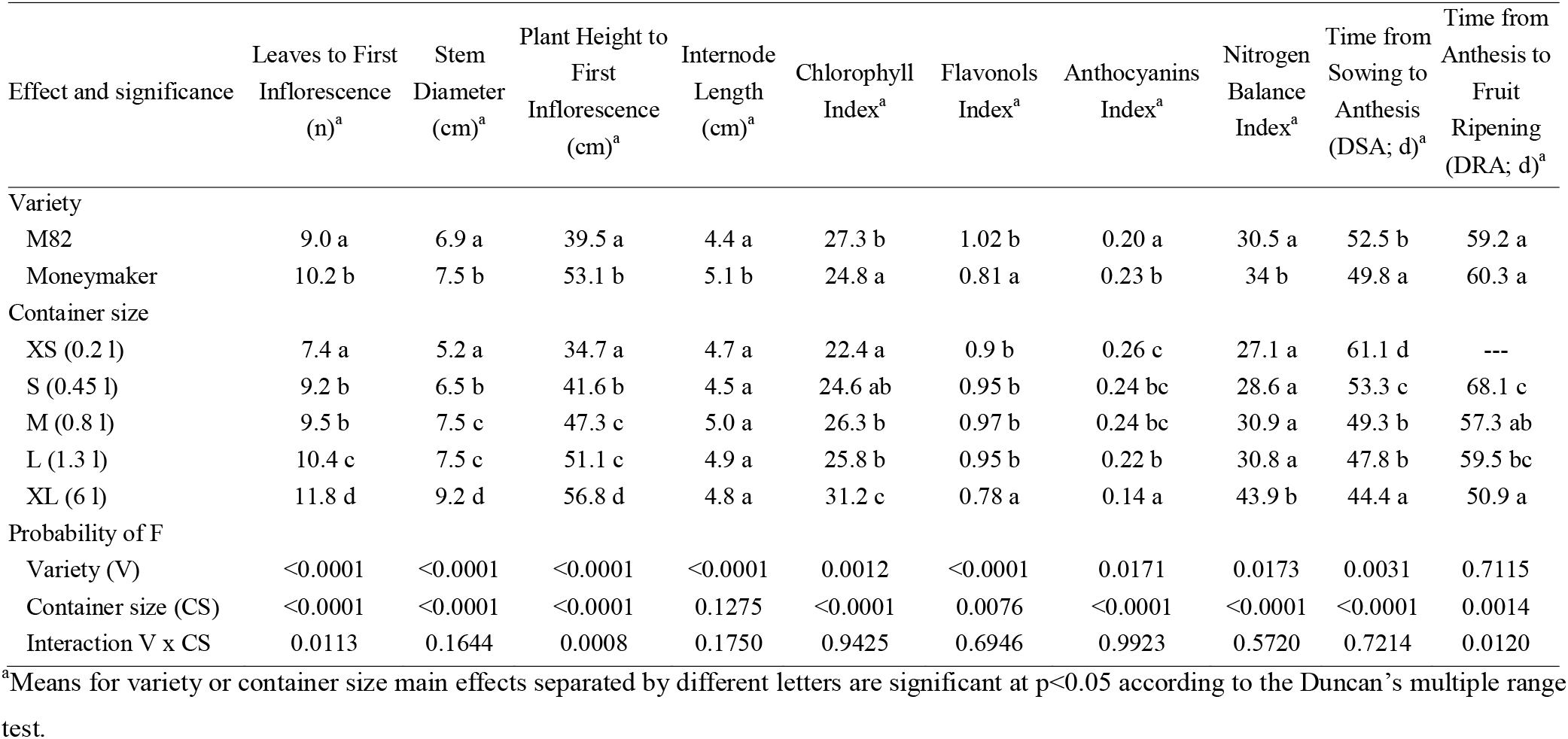
Main effects of variety and container size on growth and development, physiological and reproductive traits and significance (probability of F) of the main effects of variety and container size and their interaction in two tomato varieties grown in five container sizes (Experiment 1; EX1).

| Effect and significance | Leaves to First<br>Inflorescence<br>(n)<sup>a</sup> | Stem<br>Diameter<br>(cm)<sup>a</sup> | Plant Height to<br>First<br>Inflorescence<br>(cm)<sup>a</sup> | Internode<br>Length<br>(cm)<sup>a</sup> | Chlorophyll<br>Index<sup>a</sup> | Flavonols<br>Index<sup>a</sup> | Anthocyanins<br>Index<sup>a</sup> | Nitrogen<br>Balance<br>Index<sup>a</sup> | Time from<br>Sowing to<br>Anthesis<br>(DSA; d)<sup>a</sup> | Time from<br>Anthesis to<br>Fruit<br>Ripening<br>(DRA; d)<sup>a</sup> |
| --- | --- | --- | --- | --- | --- | --- | --- | --- | --- | --- |
| Variety |  |  |  |  |  |  |  |  |  |  |
| M82 | 9.0 a | 6.9 a | 39.5 a | 4.4 a | 27.3 b | 1.02 b | 0.20 a | 30.5 a | 52.5 b | 59.2 a |
| Moneymaker | 10.2 b | 7.5 b | 53.1 b | 5.1 b | 24.8 a | 0.81 a | 0.23 b | 34 b | 49.8 a | 60.3 a |
| Container size |  |  |  |  |  |  |  |  |  |  |
| XS (0.2 l) | 7.4 a | 5.2 a | 34.7 a | 4.7 a | 22.4 a | 0.9 b | 0.26 c | 27.1 a | 61.1 d | --- |
| S (0.45 l) | 9.2 b | 6.5 b | 41.6 b | 4.5 a | 24.6 ab | 0.95 b | 0.24 bc | 28.6 a | 53.3 c | 68.1 c |
| M (0.8 l) | 9.5 b | 7.5 c | 47.3 c | 5.0 a | 26.3 b | 0.97 b | 0.24 bc | 30.9 a | 49.3 b | 57.3 ab |
| L (1.3 l) | 10.4 c | 7.5 c | 51.1 c | 4.9 a | 25.8 b | 0.95 b | 0.22 b | 30.8 a | 47.8 b | 59.5 bc |
| XL (6 l) | 11.8 d | 9.2 d | 56.8 d | 4.8 a | 31.2 c | 0.78 a | 0.14 a | 43.9 b | 44.4 a | 50.9 a |
| Probability of F |  |  |  |  |  |  |  |  |  |  |
| Variety (V) | <0.0001 | <0.0001 | <0.0001 | <0.0001 | 0.0012 | <0.0001 | 0.0171 | 0.0173 | 0.0031 | 0.7115 |
| Container size (CS) | <0.0001 | <0.0001 | <0.0001 | 0.1275 | <0.0001 | 0.0076 | <0.0001 | <0.0001 | <0.0001 | 0.0014 |
| Interaction V x CS | 0.0113 | 0.1644 | 0.0008 | 0.1750 | 0.9425 | 0.6946 | 0.9923 | 0.5720 | 0.7214 | 0.0120 |

<sup>a</sup>Means for variety or container size main effects separated by different letters are significant at p<0.05 according to the Duncan's multiple range test.

As expected, the indeterminate Moneymaker variety had a higher average value for plant growth traits than the determinate M82. Regarding physiological traits, M82 had higher chlorophyll and flavonols indexes and lower anthocyanins and nitrogen balance indexes than Moneymaker. The time from sowing to anthesis was 2.7 d shorter in Moneymaker than in M82, while no differences were observed for the time from anthesis to fruit ripening (Table 2). Container size had a great impact on the growth and development of the plants, with more leaves to first inflorescence, larger stem diameter and plant height to the first inflorescence as the container size increased. For physiological traits, the chlorophyll index increased and the anthocyanins index decreased with container size, while the flavonols index was lower and the nitrogen balance index was higher in the XL container size compared to the other sizes (Table 2). The time from sowing to anthesis (DSA) decreased with container size, with a difference of 16.7 d in the time required for reaching anthesis for flowering between the XS and XL containers. For the time from anthesis to fruit ripening (DAR), plants from the XS container size did not produce ripe fruit with viable seeds (Table 2). However, a similar trend was observed to that found for DSA, with plants from the XL containers requiring on average 17.2 d less for DAR than those from S containers. On average, the generation time from sowing to ripe fruit for the XL containers in this autumn cycle was 95.3 d (Table 2).

Regarding significant interactions, in Moneymaker the number of leaves to the first inflorescence and plant height to the first inflorescence increased more than in M82 with container size (Figure 2). For container size in M82, DAR time in XL containers is significantly lower than in the other container sizes, while for Moneymaker, the only significant difference is between container L, which has a significantly lower DAR than container S (Figure 2).

**Figure 2.**
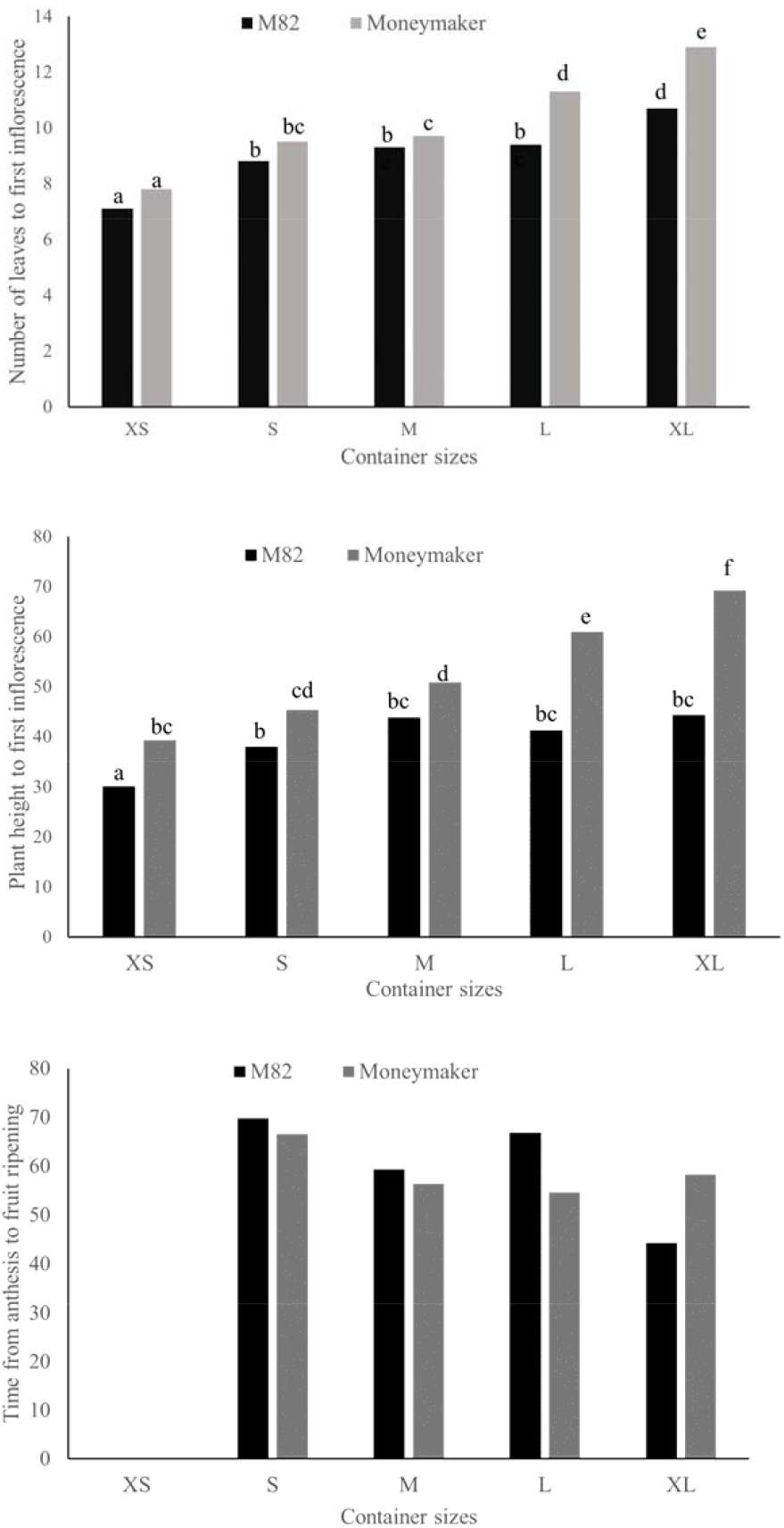
Effect of five container sizes (0.2 1, XS; 0.45 1, S; 0.8 1, M; 1.3 1, L; 6 ml, XL) on the number of leaves to first inflorescence (above), plant height to first inflorescence (center), and time from anthesis to fruit ripening (below) in M82 (black columns) and Moneymaker (grey columns) tomato plants. Means for each combination of variety and container size separated by different letters are significant at p<0.05 according to Duncan’s multiple range test.

### Effect of cold priming, water stress and nutrients supplementation on generation time (Experiment 2; EX2)

Significant (p<0.05) effects were detected for the variety (V) factor for all traits, except for the anthocyanins index (Table 3). For the treatment (T) factor, less significant differences were observed, with no significant differences for internode length and any of the four physiological traits. However, significant differences were observed for the other growth traits as well as for the times from sowing to anthesis and from anthesis to fruit ripening. The only significant V x T interaction was for the number of leaves to the first inflorescence (Table 3).

**Table 3.** Main effects of variety and treatment on growth and development, physiological and reproductive traits and significance (probability of F) of the main effects of variety and treatment and their interaction in two tomato varieties subjected to five agronomic treatments (Experiment 2; EX2).

| Effect and significance | Leaves to First Inflorescence (n)<sup>a</sup> | Stem Diameter (cm)<sup>a</sup> | Plant Height to First Inflorescence (cm)<sup>a</sup> | Internode Length (cm)<sup>a</sup> | Chlorophyll Index<sup>a</sup> | Flavonols Index<sup>a</sup> | Anthocyanins Index<sup>a</sup> | Nitrogen Balance Index<sup>a</sup> | Time from Sowing to Anthesis (DSA; d)<sup>a</sup> | Time from Anthesis to Fruit Ripening (DRA; d)<sup>a</sup> |
| --- | --- | --- | --- | --- | --- | --- | --- | --- | --- | --- |
| Variety |  |  |  |  |  |  |  |  |  |  |
| M82 | 7.2 a | 5.0 a | 35.5 a | 5.0 a | 31.2 b | 0.69 b | 0.18 a | 48.6 a | 74.5 b | 60.2 a |
| Moneymaker | 8.4 b | 6.0 b | 49.8 b | 6.0 b | 29.9 a | 0.54 a | 0.18 a | 56.5 b | 72.4 a | 65.5 b |
| Treatment |  |  |  |  |  |  |  |  |  |  |
| Control | 8.3 c | 5.7 a | 46.3 c | 5.7 a | 31.0 a | 0.63 a | 0.17 a | 52.5 a | 72.8 b | 65.5 b |
| Cold priming | 7.3 a | 5.5 a | 38.3 a | 5.3 a | 29.9 a | 0.62 a | 0.18 a | 51.3 a | 70.1 a | 62.9 b |
| Water stress | 7.5 ab | 5.3 a | 40.8 ab | 5.5 a | 31.9 a | 0.61 a | 0.18 a | 55 a | 74.1 b | 66.0 b |
| P supplementation | 8.0 bc | 5.6 a | 43.2 bc | 5.6 a | 29.6 a | 0.59 a | 0.17 a | 52.4 a | 75.3 b | 63.7 b |
| K supplementation | 7.9 bc | 5.6 a | 44.6 c | 5.6 a | 30.4 a | 0.62 a | 0.19 a | 51.6 a | 75.3 b | 56.7 a |
| Probability of F |  |  |  |  |  |  |  |  |  |  |
| Variety (V) | <0.0001 | <0.0001 | <0.0001 | <0.0001 | 0.0137 | <0.0001 | 0.9559 | <0.0001 | 0.0058 | 0.0007 |
| Treatment (T) | 0.0142 | 0.5873 | <0.0001 | 0.5873 | 0.0558 | 0.7396 | 0.9592 | 0.3175 | 0.0001 | 0.0032 |
| Interaction V x T | 0.0471 | 0.2284 | 0.0985 | 0.2284 | 0.4478 | 0.9872 | 0.4667 | 0.7169 | 0.6833 | 0.3047 |

<sup>a</sup>Means for variety or treatment main effects separated by different letters are significant at $p < 0.05$ according to the Duncan's multiple range test.

The differences observed among varieties were similar to those observed in the container size experiment (EX1), with higher average values for plant growth traits in Moneymaker than in M82 (Table 3). Similarly, for physiological traits, M82 exhibited again higher chlorophyll and flavonols indexes and lower nitrogen balance indexes than Moneymaker, although this time no differences were observed among varieties for anthocyanins index. The time from sowing to anthesis (DSA) was again shorter in Moneymaker (2.1 d) than in M82, while in contrast to EX1 the time from anthesis to fruit ripening (DAR) was shorter in M82 (5.3 d) than in Moneymaker (Table 3). The cold priming and water stress treatments reduced the number of leaves to the first inflorescence and the plant height to the first inflorescence compared to the control, while no differences were observed for internode length and any of the four physiological indexes measured. However, the cold priming treatment significantly reduced DSA with respect to the other treatments, shortening 2.7 d to the control. Also, the K supplementation treatment significantly reduced DRA compared to the other treatments, with a difference of 8.8 d with the control. On average, the generation time from sowing to ripe fruit for the control in that autumn-winter cycle was 138.3 d (Table 3).

For the only significant V x T interaction (number of leaves to the first inflorescence), no significant differences among treatments were observed for M82 while for Moneymaker the cold priming and water stress treatments the number of leaves for the cold priming and water stress treatments were lower than those of the control and K supplementation treatments (Figure 3). Also, for Moneymaker the number of leaves for the cold priming treatment was significantly lower than that of the P treatment.

**Figure 3.**
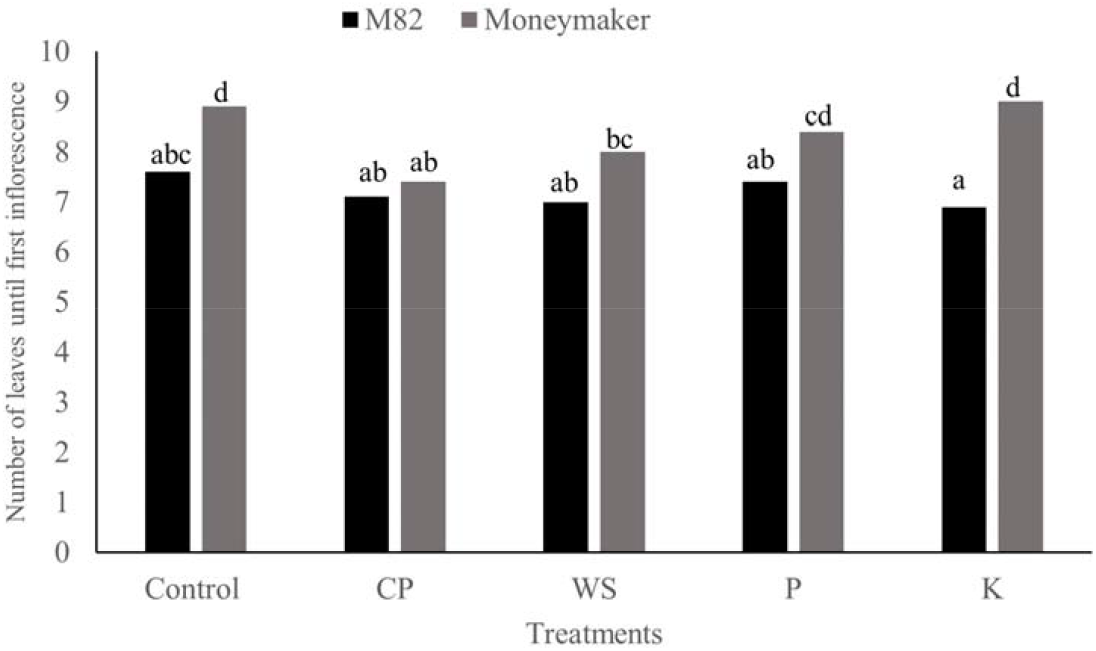
Effect of five treatments (Control, C; Cold priming, CP; Water stress, WS; P supplementation, PS; K supplementation, KS) on the number of leaves to first inflorescence in M82 (black columns) and Moneymaker (grey columns) tomato plants. Means for each combination of variety and treatment separated by different letters are significant at p<0.05 according to Duncan’s multiple range test.

### Effect of cold priming plus K supplementation and embryo rescue on generation time (Experiment 3; EX3)

Significant (p<0.05) effects were detected for the variety (V) and treatment (T) factors for the time from sowing to anthesis (DSA), time from anthesis to ripening (DRA) for plants in which fruit ripening took place in *planta*, and for the equivalent time from anthesis to ripening (eDAR) for plants in which embryo rescue was applied (Table 4). In this way, contrary to what was observed in EX1 and EX2, DSA was lower (2.2 d) in M82 than in Moneymaker. However, as occurred with the cold priming treatment in EX2, the treatment of cold priming (plus K supplementation) significantly reduced DSA (2.9 d). For the plants in which the fruits were left to ripen *in planta*, the time from anthesis to ripening (DAR) was, as in EX2, lower in M82 (4.1 d) than in Moneymaker. Also, in agreement with results from EX2 with K supplementation, the combination of cold priming plus K supplementation reduced the time from anthesis to ripening with respect to the control (3.9 d). No significant interactions V x T were observed for DRA (Table 4). On average, the generation time from sowing to ripe fruit for the control in that springer-summer cycle was 95.5 d.

**Table 4.**
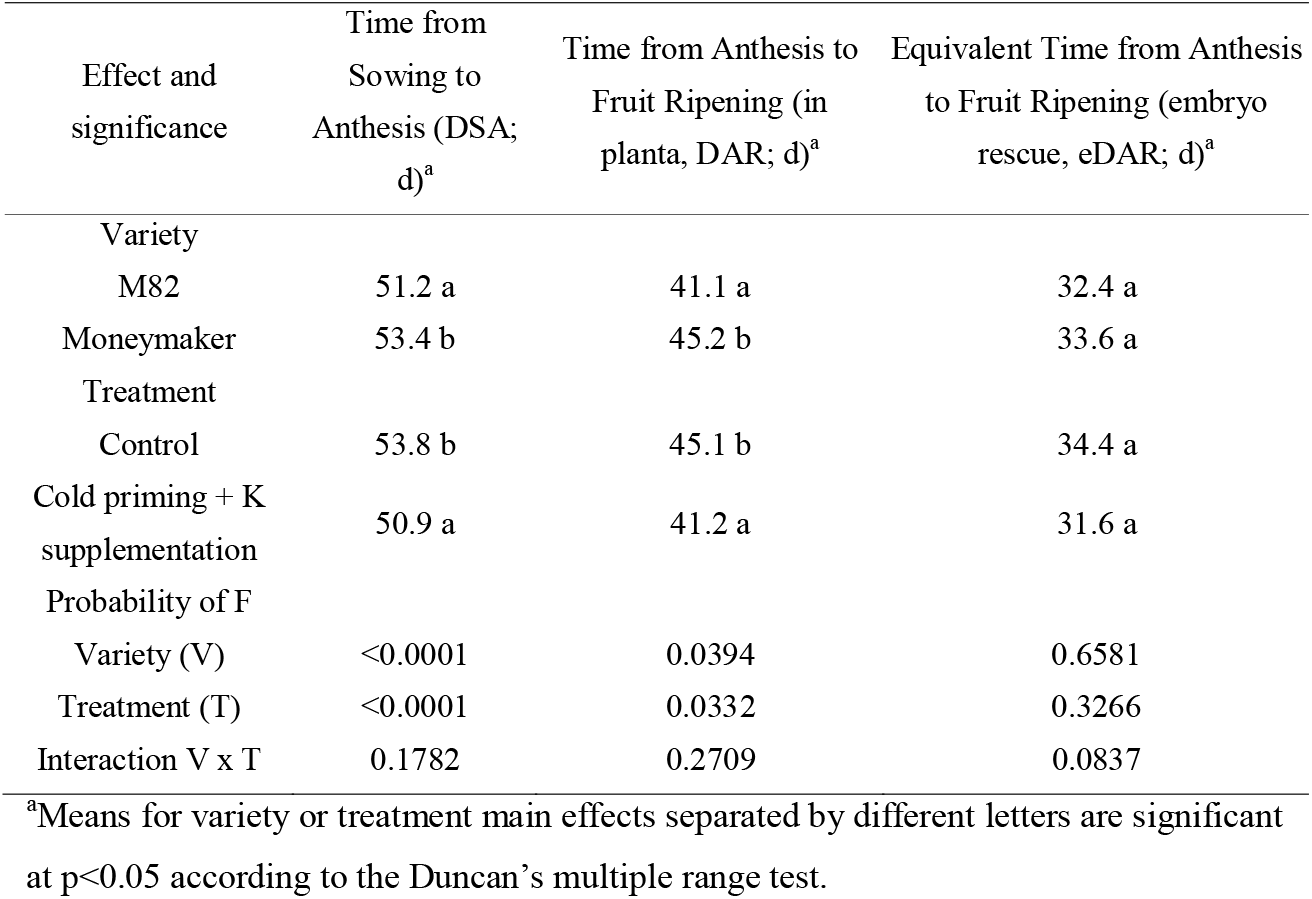
Main effects of variety and treatment on reproductive traits and significance (probability of F) of the main effects of variety and treatment and their interaction in two tomato varieties subjected to two combinations of agronomic treatments (Experiment 3). For the plants in which fruit ripening took place in planta, the number of days from anthesis to fruit ripening are presented, while for plants in which embryo rescue was practised, the equivalent time from anthesis to fruit ripening (eDAR is calculated as eDAR=DA3L-DS3L, in which DA3L is the time between anthesis and first acclimatized plant with three true leaves and DS3L is the time required from seed germination until plants with three true leaves are obtained).

Regarding plants that were left for embryo rescue, most of the embryos rescued were at the torpedo (71.7% for M82 and 44.2% for Moneymaker) or pre-cotyledonary (28.3% for M82 and 55.8% for Moneymaker) stages. A high percentage of the embryos rescued developed into plants, with 57.9% (torpedo) and 56.7% (pre-cotyledonary) embryos of M82 and 47.8% (torpedo) and 93.1% (pre-cotyledonary) embryos of Moneymaker developing plantlets that acclimatized well. To calculate the eDAR, the time required from sowing to having plantlets with three true leaves (DS3L; 30.9 d for M82 and 31.0 d for Moneymaker) was subtracted from the time elapsed between anthesis and having acclimatized plantlets from embryo rescue with three true leaves (DA3L). The eDAR values were substantially lower than those of DAR, with an average reduction of 8.7 d for M82 and 11.6 d for Moneymaker (Table 4). For eDAR no significant effects were observed for the main effects of variety (V) and treatment (T) nor for the interaction V x T (Table 4).

Given that, when using embryo rescue cold priming plus K supplementation has no effect on eDAR, we estimate that the effect of cold priming plus K supplementation with embryo rescue reduces generation time by 11.6 d in M82 and 14.5 d in Moneymaker in the spring-summer cycle (Table 4). If no embryo rescue is used (only cold priming plus K supplementation), then the reduction of generation time in this cycle would be 6.8 d in both cultivars (Table 4).

## Discussion

Tomato is the most-produced vegetable in the world (FAOSTAT, 2020) and an experimental model plant for many genetics and physiological studies (Schwarz et al., 2014). However, unlike other major crops (Ghosh et al., 2018), there is a lack of protocols to shorten tomato growing cycles. In this study, we have shown that by combining agronomic practices and embryo rescue it is possible to reduce the generation time in determinate (M82) and indeterminate (Moneymaker) tomato. The protocol we have devised, to our knowledge, is the first one combining different practices that can be easily adopted by most breeding companies and research laboratories aimed at facilitating speed breeding in tomato.

We have found that, in contrast to other crops (Zheng et al., 2013; Ferrie and Polowick, 2020), small container sizes result in a significant delay in tomato flowering and fruit ripening. While stress caused by the restriction of root growth due to small container size is known to induce flowering in some species (Takeno, 2016), in other species, including tomato (Shi et al., 2008), causes hormonal imbalances, reduction of photosynthesis and nutrient deficiencies. We have found that in tomato the reduction in growth rate coupled with a poorer physiological and nutritional status as a consequence of small container sizes results in delayed flowering and ripening. In this way, the lower values in chlorophylls and nitrogen balance index as the container size is reduced are an indicator of a suboptimal nutritional status (Farneselli et al., 2010; Cerovic et al., 2012), while the higher values of flavonols and anthocyanins indicate higher levels of stress (Kovinich et al. 2014; Martínez et al., 2016; da Silva et al., 2021). The results of the reduction of container size are similar in both varieties, although some small differences among them can be attributed to the different growth habits caused by gene variation in the SELF-PRUNING (SP) gene (Vicente et al., 2015). Our results suggest that large container sizes (6 l) are appropriate for rapid generation advancement in tomato and confirm previous results obtained by Ruff et al. (1987) who found a delay in flowering time in tomato plants grown in small pots. In another study, the use of 12 l containers was recommended as the best option for long-term evaluation experiments in tomato (Schwarz et al., 2014). However, as for rapid advancement generation only the first seeded fruit is required, we have found that 6 l containers are appropriate and allow saving space and substrate compared to larger sizes.

In the two experiments in which cold priming (alone or in combination with K supplementation) has been used, we found a reduction in the time from sowing to anthesis. It also reduced the number of leaves to the first inflorescence and the plant height to the first inflorescence. In this way, we have confirmed previous works (Lewis et al., 1953; Calvert et al., 1957; Dieleman and Heuvelink, 1992) indicating that the application of cold temperatures after the cotyledon expansion (sensitive period) reduced the number of leaves to first inflorescence and advanced flowering. In young tomato plants, the application of cold (10 °C) stress, resulted in many changes at the hormonal and gene expression levels (Zhou et al., 2019). However, to our knowledge, no works have evaluated the differential expression of genes during the sensitive period in tomato, although in *Arabidopsis* it was reported that vernalization induces the expression of FLOWERING LOCUS T (FT) which promotes flowering (He et al., 2020). Gene expression and plant growth regulators concentrations analysis during the sensitive period probably could contribute to identifying the genetic and physiological mechanisms involved in the early flowering of tomato in response to cold priming after the expansion of cotyledons.

High levels of K supplementation to the fertilizers already present in the substrate and applied as dressing fertilization resulted in a reduction of the time required from anthesis to fruit ripening in the two experiments in which it was applied, either alone or in combination with cold priming. Our results are in contrast to those observed by Besford and Maw (1975), who found an advancement in flowering at high doses of K and a faster ripening rate at low K doses. These discrepancies are probably caused by the different levels of K fertilization, which at the lowest levels applied by Besford and Maw (1975) resulted in a deficiency of K for the plant. K has an important role in tomato ripening and low levels of K availability are associated with fruit disorders (Hartz et al., 1999). Appropriate levels of K fertilization improve yield and fruit quality (Hartz et al., 2005; Caretto et al., 2008), and many metabolic changes occur in the tomato fruit as a result of different levels of K fertilization (Weinert et al., 2021). However, to our knowledge, the positive effect of K supplementation on advancing ripening time in tomato had not been reported previously. Nevertheless, Wang et al. (2021) found that at 47 d after anthesis the hue angle was lower (e.g., redder) in fruits from the high K fertilization level than those from the low K fertilization level, which could be an indication that ripening had proceeded faster in the fruits with higher K fertilization. The mechanisms involved in the faster ripening caused by K supplementation are unknown, although different levels of K supply result in changes in gene expression in multiple genes (Zhao et al., 2018), some of which may affect the ripening process.

Water stress and P supplementation did not have any significant effect on the time from sowing to anthesis or the time from anthesis to fruit ripening. Although contrasting reports exist on the effect of water stress on tomato earliness (Wudiri and Henderson, 1985; Martínez-Cuenca et al., 2020; Chong et al., 2022) the level of stress imposed is likely responsible for the differences observed. In our case, the level of water stress applied was moderate, resulting only in a reduction in the number of leaves to the first inflorescence and of the plant height to the first inflorescence, probably as a consequence of the reduced growth induced by a restriction in water availability (Gupta et al., 2020). Regarding P supplementation, it had no effect compared to the control for any of the traits evaluated. Although Dumas (1987) found that P advanced earliness in tomato, this effect was visible when compared with the non-fertilized control, which probably resulted in a suboptimal supply to the plant causing a delay in growth. Therefore, according to our results, P supplementation does not show promise for advancing generation time in tomato. The fact that none of the physiological indexes was significantly affected by the treatments indicates that, contrary to what was observed for the smaller container sizes, the plants from the different treatments grown in 6 l containers did not suffer from physiological stress (Cerovic et al., 2012). This confirms that this container size is appropriate for speed breeding in tomato.

Cold priming and K supplementation, when combined, seem to have an additive effect, with a reduction in time from sowing to anthesis and from anthesis to fruit ripening, putatively caused, respectively, by cold priming and K supplementation. This suggests that the effects of these two treatments on physiological processes, affecting both reproductive phases, are largely independent in tomato. At the phenotypic level, both traits (time from sowing to anthesis and from anthesis to fruit ripening) have been found to display a low negative correlation in a collection of 191 tomato cultivars (Wang et al., 2020), suggesting that they are largely independent. Our results also suggest that cold priming could be of interest for enhancing the earliness of commercially grown tomato, as this would just require placing nursery trays at the appropriate sensitive period (after expansion of cotyledons) in a growth chamber at low temperature for 8 d at 14°C. However, while K supplementation may be of interest for speed breeding, it does not seem appropriate for commercial sustainable tomato cultivation, given the high levels of K fertilization that would be required to reach the levels we used in our container experiments.

Embryo rescue has proved as a highly efficient tool for advancing generation time in many crops (Zheng et al., 2013; Ghosh et al., 2018; Samantara et al., 2022; Wanga et al., 2022). We have confirmed previous results revealing that embryo rescue is a powerful tool for rapid generation advancement in tomato (Bhattarai et al., 2009; Geboloğlu et al., 2011). By using embryo rescue, compared to using seeds from *in planta* ripened plants, we have found that a reduction of the growing cycle of over one week can be obtained in the spring-summer growing cycle. When combined with cold priming, which contributes to reducing the time from sowing to anthesis, the reduction of the generation time decreases by around two weeks. It is important to notice that when embryo rescue is used supplementation with K probably does not make an effective contribution to reducing the generation cycle, as our results indicate that this treatment reduces the time from anthesis to ripening only when fruits are left to ripen *in planta*. However, cold priming can be applied even to plantlets from in vitro culture at the appropriate stage (after cotyledon expansion).

Our results make a significant contribution to increasing the number of generations per year that can be normally obtained in a tomato breeding programme from the current three to almost four. Although our results represent an improvement in the number of generations per year in tomato, it is still far from the high numbers of generations that can be obtained in other crops, such as barley, in which up to nine generations per year can be obtained (Zheng et al., 2013), but are similar to important annual crops such as canola, pigeon pea or rice, in which usually four generations per year are obtained using speed breeding techniques (Wanga et al., 2021).

## Conclusions

We have found that a substantial reduction in the generation time of tomato can be achieved by a combination of agronomic techniques and embryo rescue. Stress caused by the restriction of root growth caused by small containers delayed flowering and ripening times and therefore large containers are required for fast development and shorter generation times. Cold priming and K supplementation allowed, respectively, an advancement of flowering and fruit ripening of several days. Embryo rescue at the torpedo or pre-cotyledonary stage resulted in a reduction in the generation time of several weeks. When cold priming and K supplementation in tomato plants grown in large containers are combined with embryo rescue, the average number of generations that can be obtained per year can be increased from three to almost four. The use of other complementary agronomic techniques, such as the manipulation of photoperiod, light intensity and temperatures, as well as genetic approaches may result in additional reductions in generation time in tomato.

## Funding

This work has been funded by the European Commission H2020 Research and Innovation Programme through the HARNESSTOM innovation action (grant agreement No. 101000716) and by grant CIPROM/2021/020 (project SOLECO) funded by Conselleria d’Innovació, Universitats, Ciència i Societat Digital (Generalitat Valenciana, Spain). Pietro Gramazio is grateful to Spanish Ministerio de Ciencia e Innovación for a post-doctoral grant (RYC2021-031999-I) funded by MCIN/AEI/10.13039/501100011033 and the European Union through NextGenerationEU/PRTR.

## Declaration of Competing Interest

The authors declare that they have no known competing financial interests or personal relationships that could have appeared to influence the work reported in this paper.

## Notes

### Competing Interest Statement

The authors have declared no competing interest.

